# X-ray Spectroscopy Meets Native Mass Spectrometry: Probing Gas-phase Protein Complexes

**DOI:** 10.1101/2025.02.14.638288

**Authors:** Jocky C. K. Kung, Alan Kádek, Knut Kölbel, Steffi Bandelow, Sadia Bari, Jens Buck, Carl Caleman, Jan Commandeur, Tomislav Damjanović, Simon Dörner, Karim Fahmy, Lara Flacht, Johannes Heidemann, Khon Huynh, Janine-Denise Kopicki, Boris Krichel, Julia Lockhauserbäumer, Kristina Lorenzen, Yinfei Lu, Ronja Pogan, Jasmin Rehmann, Kira Schamoni-Kast, Lucas Schwob, Lutz Schweikhard, Sebastian Springer, Pamela H.W. Svensson, Florian Simke, Florian Trinter, Sven Toleikis, Thomas Kierspel, Charlotte Uetrecht

## Abstract

Gas-phase activation and dissociation studies of biomolecules, proteins and their non-covalent complexes using X-rays hold great promise for revealing new insights into the structure and function of biological samples. This is due to the unique properties of X-ray molecular interactions, such as site-specific and rapid ionization. In this perspective, we report and discuss the promise of first proof-of-principle studies of X-ray-induced dissociation of native biological samples ranging from small 17 kDa monomeric proteins up to large 808 kDa non-covalent protein assemblies conducted at a synchrotron (PETRA III) and a free-electron laser (FLASH2). A commercially available quadrupole time-of-flight mass spectrometer (Q-ToF2, Micromass/Waters), modified for high-mass analysis by MS Vision, was further adapted for integration with the open ports at the corresponding beamlines. The protein complexes were transferred natively into the gas phase via nano-electrospray ionization and subsequently probed by extreme ultraviolet (FLASH2) or soft X-ray (PETRA III) radiation, in either their folded state or following collision-induced activation in the gas phase. Depending on the size of the biomolecule and the activation method, protein fragmentation, dissociation, or enhanced ionization were observed. Additionally, an extension of the setup by ion mobility is described, which can serve as a powerful tool for structural separation of biomolecules prior to X-ray probing. The first experimental results are discussed in the broader context of current and upcoming X-ray sources, highlighting their potential for advancing structural biology in the future.

## Introduction

Electrospray ionization (ESI) of proteins and their complexes in combination with mass spectrometry (MS) is nowadays a standard technique in biophysics and structural biology due to its ease of application and its versatility of experiments. MS has been particularly successful in probing two fundamental aspects of proteins: their structure and sequence. Structural analysis in the form of conformational studies of biomolecules has been largely enabled by advances in native MS,^1^ which gently transfers proteins and their complexes from native-like aqueous solutions into the gas phase under close to native conditions (usually by nano-ESI^2^). This has been proven numerous times indirectly with ion mobility (IM) measurements,^3^ free-electron laser (FEL) spectroscopy^,4^ and through direct observation of single molecules via electron microscopy imaging subsequent to soft-landing of natively spayed proteins^.5^

Tandem MS, where ions are isolated by their mass-to-charge ratio (*m/z*) and analyzed after being subjected to fragmentation, is extremely powerful for both sequence and structure analysis. Laboratory-based fragmentation methods include collision-induced dissociation (CID), electron-based dissociation (ExD),^6^ surface-induced dissociation (SID)^,7^ or infrared/ultraviolet photodissociation (IRMPD/UVPD)^.8,9^ Each method of fragmentation has its own strengths and weaknesses. CID and IRMPD require cycles of slow heating of the sample by collisions with background gas and intramolecular vibrational redistribution (IVR). UVPD/ExD are faster fragmentation methods and can therefore be more sensitive to the initial conformation of the samples^.10^

Similarly, X-ray excitation induced fragmentation of proteins may offer advantages such as site specificity and speed. A single absorbed photon is usually followed by an ultrafast Auger–Meitner decay (~fs), resulting in rapid energy deposition and potentially structure-dependent fragmentation. Studies of the physical process of the excitation of biomolecules by X-rays, such as peptides and proteins, have been conducted previously^.11–14^ The mechanism of relaxation and the fragmentation pathways are heavily dependent on the size of the molecule. For small molecules, ejection of hydrogens or protons and small fragments are dominant. For smaller peptides, core-electron photoionization and subsequent relaxation processes (both the emission of photoelectrons and Auger–Meitner electrons) are followed by fragmentation pathways that produce small *m/z* fragments.^11^ For larger proteins, IVR of energy can outcompete the fragmentation pathways, reducing the number of fragments produced after ionization.

Using X-rays for the study of protein complexes may provide unique and complementary information that is not present in other structural-biology techniques. However, the interaction of large gaseous protein complexes, made feasible by native MS, with X-rays is yet not well understood. In order to take advantage of the rapid rate of relaxation by fragmentation as a tool to study the conformation of large biomolecules, it is necessary to understand the underlying physics of the process.

Moreover, X-ray fragmentation may also be a complementary technique in (native) top-down (TD) MS, where protein sequence coverage can be achieved via fragmentation of the whole protein in the mass spectrometer.^15,16^ The fast rate and rapid energy deposition after the absorption of an X-ray photon is of particular interest in TDMS, as X-ray photodissociation may become a complementary method for better protein sequence coverage and survival of post-translational modification. In addition, X-ray fragmentation can be combined with other established gas-phase techniques such as ion-mobility spectrometry (IMS). In combination with simulations, IM-tandem MS can be a powerful tool for structural analysis of protein complexes.^17,18^

Furthermore, native MS can be used to study radiation damage in biomolecules in the gas phase, focusing on secondary ionization events from photo- and Auger–Meitner electrons—similar to those occurring in X-ray radiation-induced damage to biological tissues.^19^ A better understanding of the effects of X-rays on biomolecules, including proteins, is crucial in the biomedical field. In this isolated gas-phase environment, native MS enables the study of X-ray effects on proteins without interference from environmental and surrounding factors.

Here, we present the first proof-of-principle experiments of our campaign to study the interaction of X-rays with gaseous proteins and their non-covalent complexes of various sizes obtained by native MS. For our experiments, a commercially available high-mass Q-ToF mass spectrometer further modified for X-ray experiments was installed at multiple open-port beamlines at DESY in Hamburg, Germany. We focus on data collected from two beamlines, the synchrotron beamline P04 at PETRA III^,20^ as well as the FL24 beamline of the free-electron laser FLASH2^21^. The experimental MS setup is capable of measuring proteins and protein complexes ranging from small peptides up to MDa virus-like particles (VLPs) with a diameter of more than ~30-40 nm^.22–24^ In addition to a quadrupole mass filter (QMF), which was used to select molecular ions of a specific *m/z* in the gas phase before the X-ray interaction region, we demonstrate the use of CID and IM prior to probing the proteins by X-rays for pre-activation and conformational separation, respectively. The presented results are discussed in the context of future potential experiments and their current limitations.

### Instrumental setup and experimental methods

Fig. 1 shows a schematic drawing of an X-ray coupled version of the quadrupole time-of-flight (Q-ToF2, Micromass/ MS Vision) spectrometer modified for high mass.^25^ Modifications beyond high-*m/z* capability include open-port access via a DN40 ConFlat (CF) vacuum flange at the transfer hexapole ion guide behind the collision cell for coupling to P04^20^ at the PETRA III synchrotron or FL24 at the FLASH2 FEL^.26,27^ The full details of the modifications for optical access are described in the Supplementary information.

**Fig. 1.**
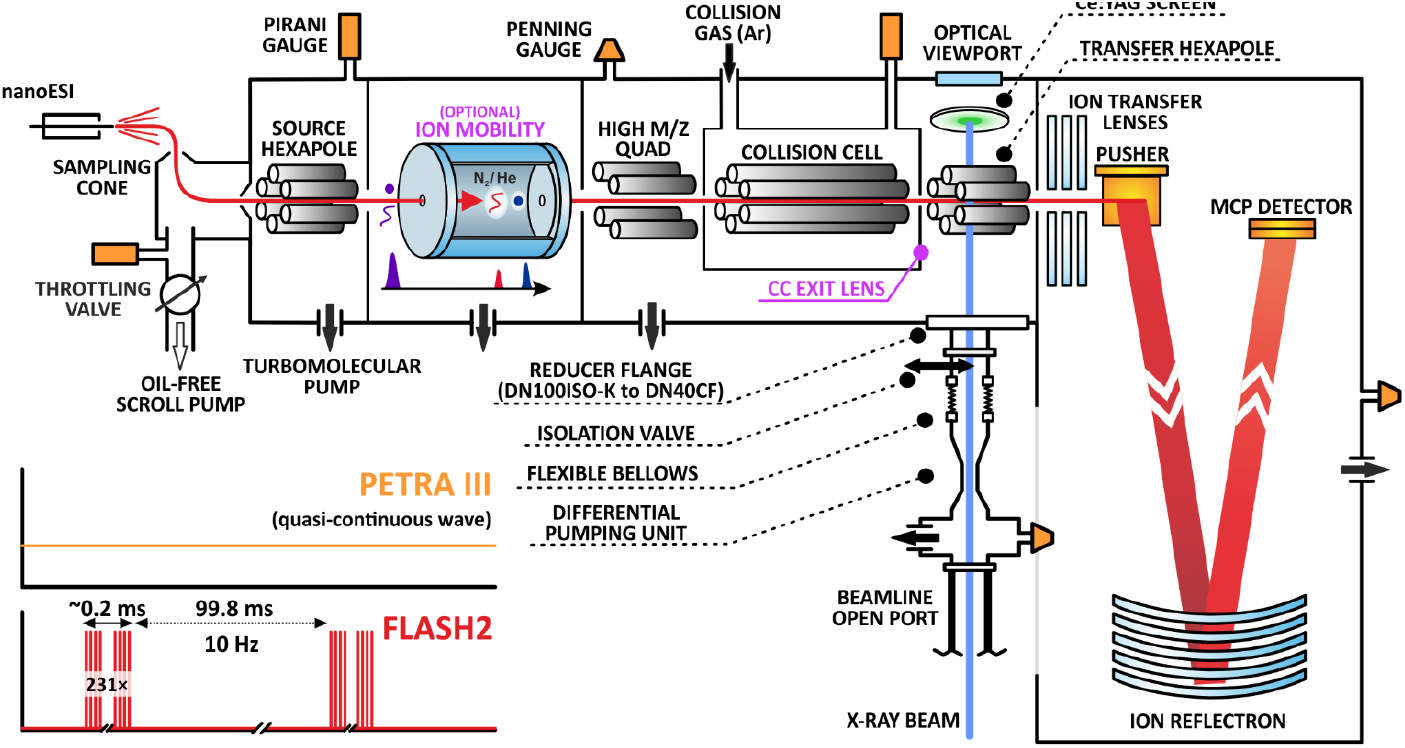
Schematic of the experimental setup. Ions are generated from solution in a nano-ESI source into the gas-phase and transported through a high-mass modified Q-ToF mass spectrometer to a microchannel plate (MCP) detector. As they fly past, they intersect perpendicularly with X-ray / EUV photons in a radially confining hexapole between the collision cell and the ToF analyzer. An optical viewport and a Ce:YAG screen were used to aid the alignment of the setup with respect to the photon beam. Magenta labels show the optional ion mobility device used for conformational separation and the collision cell (CC) exit lens used to temporarily trap ions as described in the main text. The inset illustrates the photon delivery structure of both the quasi-continuous PETRA III synchrotron (equally distributed pulses with 16 ns or 192 ns microbunch spacing when operated at 62.5 MHz and 5.2 MHz, respectively) and the unevenly pulsed FLASH2 free-electron laser used for the reported experiments.

For the experimental sequence, samples are transferred into the gas phase using ESI or nano-ESI. All protein and peptide samples or other materials are described in the Supplementary information. Experiments were performed in positive ion mode. Hence, ESI produces positively charged protein ions that enter the instrument. Afterwards, the ions were selected based on their *m/z* using the QMF of the Q-ToF. Depending on the pressure in, and the voltage gradient across the collision cell, samples were either merely thermalized and transported through the collision cell, which is aided by gentle collisional cooling and beam focusing, or vibrationally activated in the gas phase via energetic collisions with argon, before they were transported to the transfer hexapole ion guide and probed perpendicularly by the X-rays. The X-ray beam was typically much smaller in diameter—at most one-tenth the size of the ion beam— allowing only a few percent of the direct ion beam to be probed by the X-rays. Ions (including the products) were then transferred to the pusher region, mass analyzed in the ToF and detected upon hitting a microchannel plate. Additional instrument components and measurement steps for pulsed operation were required at FLASH2 due to the long time period between the photon bunches, see the Supplementary information for further details.

For the ion-mobility X-ray experiments only, an additional vacuum chamber containing a resistive glass drift tube was installed in front of the source hexapole of the Q-ToF. The instrument components were obtained from MS Vision and the design is based on the MoQToF instrument by Barran and co-workers.^28,29^

## Results and discussion

### Fragmentation

Protein complex fragmentation experiments were performed at two different X-ray light sources, each corresponding to a distinct ionization regime. At the P04 beamline of PETRA III, protein complexes were probed via single-photon 1s core-ionization at a photon energy of 595 eV using a pink (polychromatic) beam. In contrast, at the FL24 beamline of FLASH2, protein samples were probed via multi-photon inner-shell ionization at a photon energy of 163 eV and a pulse energy of 140 µJ, with a focus diameter (full width at half maximum) of 100 µm. Here, the absolute number of absorbed photons is exceedingly difficult to estimate due to the complexity of the underlying ionization processes. For example, in the case of haemoglobin (Hb, Fig. 2 d), these FEL parameters suggest the absorption of up to a couple hundred photons per FEL pulse. This estimate is based on the independent atom model, and neglects, e.g. changes in absorption cross section due to the molecular orbitals, or the increased ionization of the protein complex. In turn each photo-electron creates up to 7 secondary electrons, due to electron impact ionization, within a volume with a radii of around 2.5 nm.^30^ The resulting number necessitates a very high state of ionization.

**Fig. 2.**
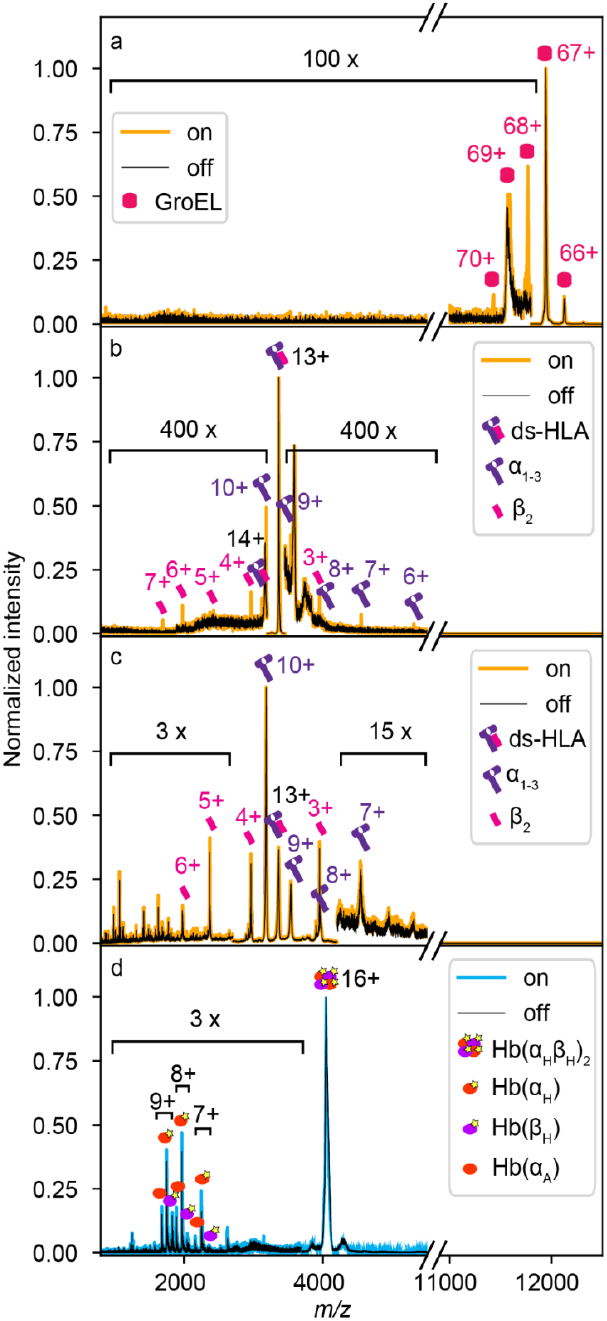
Mass spectra of protein complexes in the presence (on, yellow or blue) and absence (off, black) of X-ray excitation. Spectra in panels a-c were measured at the PETRA III P04 beamline. Spectra in panel a show non-collisionally activated GroEL. In panels b and c, mass spectra of non-collisionally activated and collisionally activated disulfide-stabilized human leukocyte antigen (ds-HLA), respectively, are depicted. Spectra of collisionally activated tetrameric haemoglobin (Hb) in panel d were measured at the FL24 beamline of the FLASH2 FEL.

Figure 2 shows a selection of protein complex mass spectra recorded with the instrumental setup at the two facilities. The spectra compare the protein complexes in the presence (on) and absence (off, black) of X-ray irradiation. Spectra in Figs. 2a-c were measured at the P04 beamline (yellow), the spectra in Fig. 2d were measured at the FLASH2 facility (blue). Additionally in Fig. 2a and 2b, non-preactivated, native-like protein complexes, of the bacterial chaperone GroEL (≈ 808 kDa, 14-mer) and disulfide-stabilized human leukocyte antigen (ds-HLA, ≈ 44 kDa, heterodimer comprising α_1-3_ heavy chain and β_2_-microglobulin) are shown.^31^ In contrast, Fig. 2c and 2d depict spectra of collisionally pre-activated ds-HLA and human haemoglobin (Hb, ≈ 64 kDa, heterotetramer comprising two α and two β subunits with or without a haem cofactor - holo-/apo-, respectively) protein samples, i.e., non-native and partially fragmented protein complexes. Labelled peaks show the assigned, highlighted fragments as well as the corresponding charge states.

Generally, the measured X-ray induced mass spectra are dominated by the direct ion beam, which can be attributed to the previously mentioned mismatch in ion and X-ray beam diameters, as well as the low probability of X-ray interaction as the ions were irradiated on-the-flight without extended ion trapping. The collisionally pre-activated samples in Fig. 2c-d exhibit a much higher ion yield, as evidenced by the significantly lower magnification factors needed to visualize the X-ray-induced fragmentation channels. This can be qualitatively explained by the higher internal vibrational energy of the samples due to CID, which results in a lower energy threshold required for fragmentation. Moreover, contribution of a difference in secondary ionization events due to a change of conformation cannot be excluded either.

Fig. 2a shows the quadrupole-filtered and non-activated (native-like folded) GroEL. In the ‘X-ray off’ spectrum (black), the precursor ions of the 67+ charge state dominate the spectrum. Small amounts of 66+ and 69+ ions are present due to charge stripping of the complex accompanied by additional desolvation required after isolation in the QMF and/or incomplete mass filtering by the quadrupole itself. But their relative intensity is below 1% compared to 67+ and negligible for the X-ray interaction. Upon X-ray irradiation, the 67+ charge state of the very large oligomer did not fragment, but multiple ionization events occurred, which can be explained by photoelectron and Auger–Meitner electron relaxation mechanisms. The lack of fragmentation can be attributed to the size of GroEL, as larger complexes have more vibrational degrees of freedom to absorb the excitation energy, which is in line with studies conducted by Schlathölter and co-workers.^11^

Fig. 2b shows a similar initial situation for the non-activated heterodimer ds-HLA. The 13+ charge state is quadrupole-filtered and dominates the spectrum. In contrast to GroEL, the protein complex also shows a relatively minor peak for secondary ionization to 14+ but in addition primarily undergoes fragmentation into its α_1-3_ and β_2_ subunits, similar to the products in an SID experiment.^32^ The summed final charge states of the products can be estimated based on the relative peak intensities of the product ion peaks, which match ions with 13+, 14+, and 15+ charge states.

In the case of QMF and collisionally pre-activated ds-HLA (Fig. 2 c), the situation is more complex. The spectrum is dominated by the α_1-3_ 10+ charge state, a fragment generated during the CID process. The initial quadrupole-filtered 13+ charge state has an intensity comparable to that of other ions produced through CID fragmentation. Thus, the X-rays interact with multiple ionic species at once. A general increase of the already populated fragmentation channels is visible, and new fragmentation channels could be hidden in the significantly increased ion background due to the CID process. However, the ion signal of the β_2_ 3+, 4+, 5+, and 6+ subunits are clearly enhanced upon X-ray interaction, suggesting an origin in the ds-HLA 13+ ions, as these subunits cannot come from the α_1-3_ 10+ charge state. This pattern resembles that in Fig. 2b but occurs with significantly higher fragmentation efficiency. It can be explained by the increased internal energy of the protein complex due to the collisional activation prior to X-ray exposure.

Fig. 2d shows mass spectra of collisionally activated and quadrupole-filtered Hb 16+ proteins. In contrast to Fig. 2c, the CID spectrum is dominated by a single species. CID fragments include both holo- (haem bound) and apo- (without haem) forms of monomers of both subunits, with intensities ranging from 1 to 20% of the main peak. Similar to Fig. 2c, the ionization due to the FEL primarily enhances existing CID fragmentation channels at a comparable high yield, albeit with slightly different branching ratios. This similarity suggests that the detected fragments from FEL ionization primarily originated from protein complexes that were probed at the edge of the FEL focus and ionized by only a few XUV photons. As mentioned above, for Hb, a first approximation would estimate the absorption of a couple hundred photons in the focus of the FEL. Apparently, the chosen experimental setup is not optimal for this type of FEL experiment. The high number of absorbed photons likely induce a strong Coulomb explosion of the protein sample, generating small ionic fragments that are too fast to be retained and effectively transferred by the transfer hexapole with confining rf-frequencies primarily optimized for larger species. This, together with the geometric distance between the interaction zone and the ToF entrance, along with the ion spectrometer’s lower detection limit of 150 *m/z* due to overwhelming electronic signal from the pusher, prevents the detection of these fragments.

### Addition of mobility separation

Gas-phase structural techniques, such as IMS, are well established and can significantly enhance X-ray protein studies. By separating conformations, these techniques offer additional information beyond what is obtainable from X-ray fragmentation alone. Furthermore, when combined with simulations, the measured collision cross sections offer new insights for the interpretation of the resulting fragments.

Fig. 3 shows the first proof-of-principle results from IM experiments conducted with X-rays using synchrotron radiation. To enable these measurements, a custom-built IM drift tube^33^ was installed on the Q-ToF2 as shown in Fig. 3. The sample was a helix-turn-helix peptide (HTH, see Supplementary information for the sequence), similar to a study by Jarrold and co-workers^34^. The polypeptide sample was first subjected to IM, then QMF for the doubly ionized HTH peak at 1321 *m/z* before being probed by the X-rays.

**Fig. 3.**
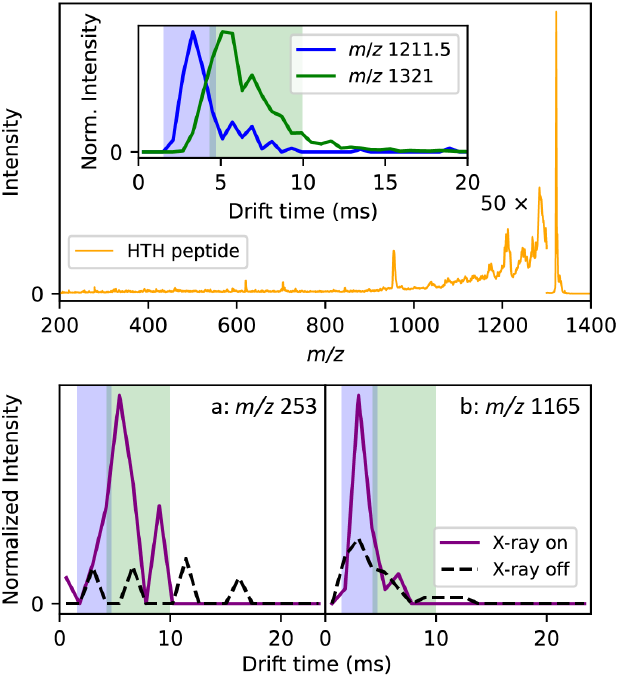
Single photon X-ray excitation of the helix-turn-helix peptide (HTH) sample. The upper panel contains the mass spectrum of the peptide after irradiation. The inset shows the arrival time distributions of the *m/z* representing HTH (*m/z* 1321) and the second most prominent peak in the mass spectrum from the sample (*m/z* 1211.5). The lower panel contains arrival time distributions of two fragment ions with and without irradiation.

As seen in the spectrum in Fig. 3 and indicated by the magnification factor, the filtering was less effective than in Fig. 2, resulting in a very low signal-to-noise ratio for this experiment. However, the ion drift time could be used to discriminate between different species. The mass spectrum of the peptide sample shows incomplete synthesis containing multiple shorter peptides. In the top panel of Fig. 3, 1321 is the expected *m/z* of the HTH peptide while 1211.5 corresponds to the *m/z* of the peptide fragment missing the last two residues. Their arrival time distribution is shown in the inset. Their drift times are different enough that fragments produced after irradiation from either the full peptide or the impurity can be distinguished; the ion-mobility separation is performed before the Q-ToF analysis, and before fragmentation. In the bottom panel, the arrival time distributions of two potential fragment ions with *m/*z 253 and 1165 with and without X-ray excitation are depicted. The blue and green shaded areas mark the arrival time peaks of the full and partial peptide of the inset in the top panel respectively.

The IM module is being further optimized, including the resolution of the measurements. Nonetheless, the distributions show the ability of distinguishing fragment ions as either being from the peptide or from impurities. As the instrument is optimized, selection of specific protein conformations before fragmentation will be feasible. This will allow the differentiation of fragmentation pathways of multiple conformers, and provide structural and sequence information as reported from other IMS combined with photodissociation techniques^.35–37^

### Summary and outlook

In these first experiments of the X-ray excitation of gaseous protein and protein complex ions, we have demonstrated the utility of native MS as a delivery system for X-ray excitation of isolated proteins, especially larger complexes in the range of 50 kDa to 1 MDa.

The fragmentation pattern and pathway after irradiation are similar to the trend shown in previous studies of peptides and smaller proteins, with additional pathways that are not possible for monomeric proteins. In the presence of only X-ray irradiation, smaller protein complexes proceed in dissociation by breaking off non-covalent interactions after induced ionization, resulting in formation of protein subunits and ligands. For very large complexes, ionization by simple electron ejection and subsequent Auger–Meitner decay dominate. When comparing, the fragment abundance from X-ray excitation is lower than in CID, although this is possibly due to low interaction probability of photons with complexes. In the present examples, X-ray excitation of already collisionally activated complexes generally enhances the CID fragmentation channels. These observations are in line with the model of vibration redistribution of energy after ejection of electrons;^11^ in which larger protein complexes have more vibrational modes to distribute this energy, unless these are already diminished by collisional activation. The addition of IMS before X-ray fragmentation is feasible and can be incorporated in the future for the study of large protein complexes.

From these initial experiments, we have recognized some limitations of the current instrumental setup and method. The major issue is the low signal-to-noise ratio. To increase this, we identified the source of the background ion signal as problematic with collisional pre-activation. That being said, QMF without pre-activation results in very low ion background, allowing high sensitivity for the low abundant products. Such an approach is suited when the subject of interest is in fragmentation mechanisms and the corresponding irradiation induced ionization of non-covalent complexes. Thus, even with the current background, native MS X-ray experiments are useful for spectroscopic and radiation damage studies.

For the increase in signal of the fragment ions, the issue is either low statistical probability of ion-photon interactions, or the difficulty in detecting product ions with high kinetic energy. As a side note, we had high dissociation yields for two samples in one campaign suggesting that higher efficiency could be achieved (Supplementary Fig. S1). Thus, improvement in the instrumental setup illustrated in Fig. 1 must be considered. We note that the photons are transmitted through the rods of the last transfer-hexapole in front of the ToF analyzer, and ions are irradiated perpendicularly, 10 cm before they enter the ion transfer lens and pusher region of the spectrometer. The location was already chosen (mechanically) as close as possible to the entrance to the ToF region. It is unclear how many ions are lost in the hexapole after X-ray interaction due to high kinetic energies obtained during the relaxation process. This concern is especially important for experiments at FEL beamlines (Fig. 2d), where strong Coulomb explosions are expected. As such, FEL (or multiphoton) experiments would profit from an interaction point in the pusher region to extract the fragments as it could for instance be realized in the MS SPIDOC setup,^29^ or a gas-filled ion trap for cooling and trapping the fragments.^11,38,39^

Another possibility to consider is that the number of interaction events between the photons and complexes was low. If the absorption cross section is high, and yet for example, the X-ray-ion beam overlap is low, then absorption rarely takes place. This can be improved by irradiation in parts of the instrument where the ions are in higher density, such as in an ion trap, or by instrumentation that allows co-axial ion and photon beams for interaction.^40–42^

Finally, in order for the fragmentation to proceed, the fragmentation after absorption of the X-ray photon is due to energy left behind by the ionization/decay processes. In larger protein complexes, the reabsorption of any photoelectrons or Auger–Meitner electrons and subsequent ionization are expected to be substantial.^43^ Therefore, the energy of the ejected electron from the core is an important parameter to investigate. This can be measured by tuning the photon energy and conducting spectroscopy experiments to gain a deeper understanding of the relaxation pathways of proteins of different sizes, following X-ray photon absorption at different energies, which is a unique feature available for these wavelengths exclusively at synchrotrons and FELs.^13^

In comparison to other MS fragmentation techniques,^15^ such as ExD or UVPD, not as many different pathways of fragmentation have appeared after X-ray irradiation. Two main strategies can be employed to use X-rays as a complementary technique for TDMS and spectroscopic experiments. The first approach involves investigating changes in X-ray photon energy, as mentioned above. The second involves MS^n^ experiments, where various activation methods (including X-rays) are combined with additional filtering of individual species after CID or X-ray interaction. This could be achieved, for instance, using an Omnitrap platform^44^, which is particularly well-suited for cycling and filtering reaction products.

In terms of the future for employing native MS as a sample delivery system for X-ray experiments, the delivery of natively folded, mass and conformationally selected protein complexes is mature for fragmentation experiments, and moreover a wide variety of other X-ray experiments of large biomolecules. The MS SPIDOC project was conceived to leverage the techniques known in mass spectrometry for background free, highly selective single particle imaging.^29,33^

The absorption of large numbers of photons, which might have been a concern for a fragmentation experiment inside a modified commercial instrument, can become an advantage for ion imaging experiments such as velocity map imaging (VMI) or Coulomb explosion imaging of biological structures ^45,46^.

Moreover, since experiments are often conducted at large light source facilities, additional lasers are available for pump-probe experiments. This enhances the potential of using native MS as a promising sample delivery method for studying structural changes in biomolecules in real time. Recent work highlights the capability of native MS to look at complex kinetics in proteins and protein complexes.^47–50^

With our initial experiments and studies, the combination of native MS and X-ray sources promises to become an invaluable tool in structural biology, biophysics and spectroscopy.

## Author contribution

**Conceptualization** – A. K., K. K., J. C., T. K., C. U. **Methodology** – J. C. K. K, A. K., K. K., St. B., J. C., T. D., Y. L., T. K., C. U. **Software** – J. C. K. K., T. D., S. D., Y. L., F. S., T. K., **Validation** – J. C. K. K., A. K., K. K., Y. L., T. K., **Formal Analysis** – J. C. K. K., A. K., K. K., Y. L., T. K., **Investigation** – J. C. K. K., A. K., K. K., St. B., Sa. B., J. B., J. C., T. D., S. D., L. F., J. H., K. H., J.-D. K., B. K., J. L., Y. L., R. P., J. R., K. S.-K., Lucas. S., P. H.W. S., F. S., F. T., S. T., T. K., C. U. **Resources** – J. B., J. C., K. F., K. L., Lutz S., S. S., F. T., S. T., C. U. **Data Curation** – K. K., T. D., Y. L., T. K., **Writing – Original Draft** – J. C. K. K., A. K., T. K., **Writing – Review & Editing** – K. K., St. B., Sa. B., J. B., C. C., S. D., L. F., J. H., K. H., J.-D. K. B. K., J. L., Y. L., R. P., J. R., K. S.-K., Lucas. S., Lutz S., S. S., P. H. W. S., F. S., F. T., S. T., C. U. **Visualization** – J. C. K. K., A. K. K. K., T. K., **Supervision** – A. K., C. C., K. L., Lucas. S., Lutz S., T. K., C. U. **Project Administration** – A. K. T. K., C. U. **Funding Acquisition** – Sa. B., C. C., Lutz S., C. U.

## Acknowledgement

We would like to thank everyone that has been involved in this experimental campaign. We acknowledge DESY (Hamburg, Germany), a member of the Helmholtz Association HGF, for the provision of experimental facilities. Parts of this research were carried out at FLASH and PETRA III. We would like to thank Moritz Hoesch and Marion Kuhlmann for assistance in using P04 and FL24 respectively. Beamtimes were allocated for proposals F-20150009, F-20171103, I-20180927, I-20190534, I-20221192. We also acknowledge members of the MS SPIDOC consortium, the Uetrecht group (Hao Yan) and collaborators, including at the Universität Greifswald (Paul Fischer, Gerrit Marx), the University of Hamburg (Henning Tidow), European XFEL (Joachim Schulz, Carsten Deiter, James Moore, Rita Graceffa, Matthäus Kitel) for their participation in the beamline experiments. Finally, we acknowledge all the funding agencies that made this project possible: ERC SPOCk’S MS funded by the ERC (grant agreement No. 759661), PIER Ideenfonds from DESY and the University of Hamburg, and MS SPIDOC funded by the European Union’s Horizon 2020 research and innovation program (grant agreement No. 801406). Leibniz Institute for Experimental Virology is supported by the Free and Hanseatic City of Hamburg and the German Federal Ministry of Health. J. C. K. K., C. C., T. D. and C. U. acknowledge support from a Röntgen Ångström Cluster grant provided by the Swedish Research Council and the Bundesministerium für Bildung und Forschung (2021-05988, 05K22PSA). A. K. gratefully acknowledges an Alexander von Humboldt postdoctoral fellowship. C. U., A. K., and Lutz S. further acknowledge funding through Bundesministerium für Bildung und Forschung (BMBF) 05K2016 VISAVIX (05K16HG1, 05K16BH1). Sa. B. and Lucas S. acknowledge funding by the Helmholtz Initiative and Networking Fund and Sa. B. acknowledges further support by the Cluster of Excellence ‘CUI: Advanced Imaging of Matter’ of the Deutsche Forschungsgemeinschaft (DFG) – EXC 2056 – project ID 390715994. C. C. and P. H. W. S. acknowledge support from the Swedish Research Council through projects 2021-05988 and 2018-00740. C. C. acknowledges support from the Helmholtz Association through the Center for Free-Electron Laser Science at DESY. S. D. acknowledges funding from VirMScan, the Bundesministerium für Bildung und Forschung (BMBF 13GW0622). L. F. was funded by the Promotion of young CSSB scientists’ program by the Joachim Herz Foundation. F. T. acknowledges funding by the Deutsche Forschungsgemeinschaft (DFG, German Research Foundation) - Project 509471550, Emmy Noether Programme.

