## Supplementary information for "X-ray Spectroscopy Meets Native Mass Spectrometry: Probing Gas-phase Protein Complexes"

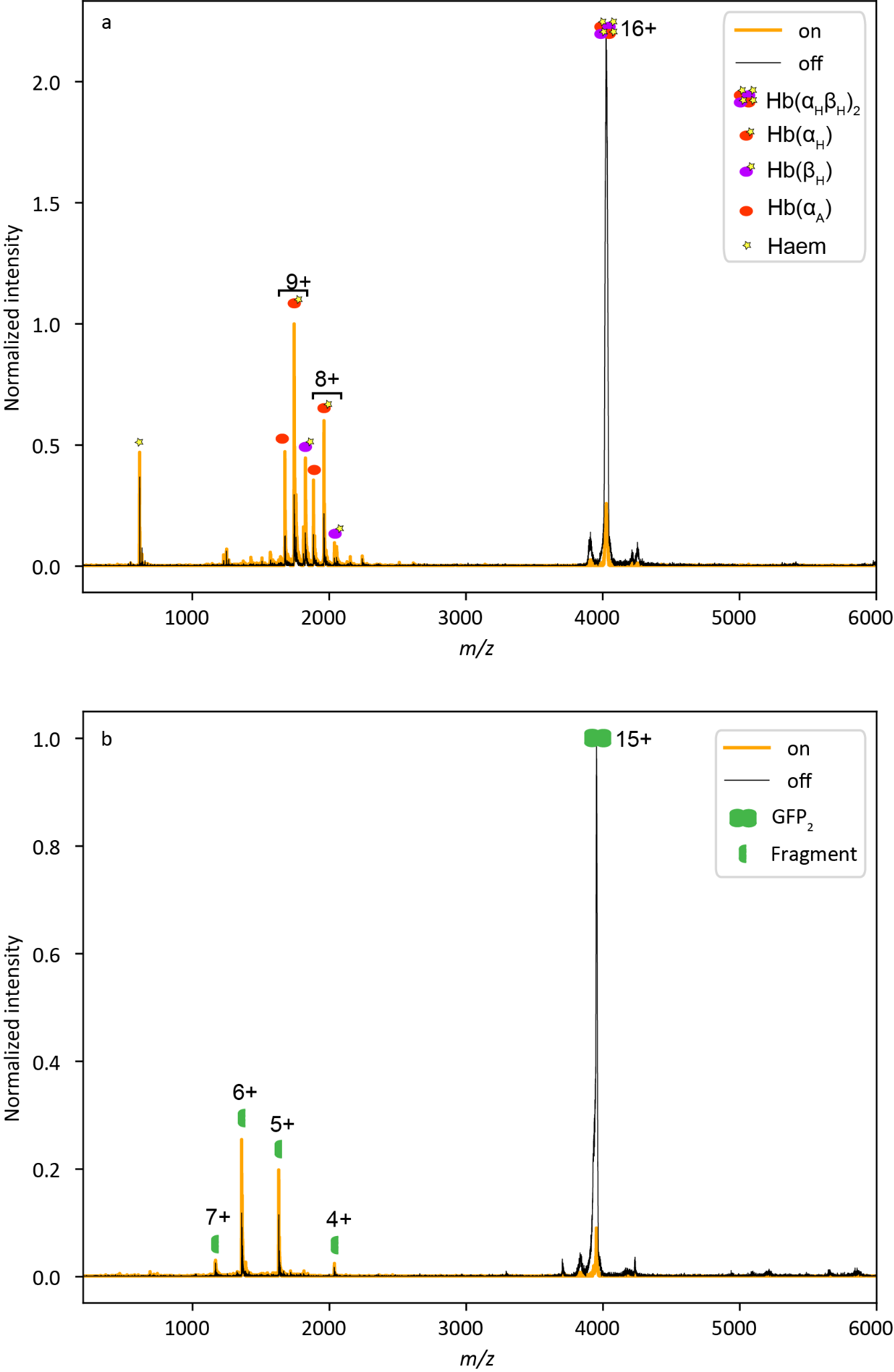

**Fig. S1** Mass spectra of haemoglobin (a) and GFP mutant (b) in the presence (on) and absence (off) of X-ray excitation during PETRA III April 2018 experiment.

**Materials**

Standard native MS protein complexes used in this study were obtained from Sigma-Aldrich (with their product number in brackets) human haemoglobin (#H7379) and chaperonin GroEL from E. coli (#C7688). GroEL was additionally repurified by precipitation as described before^1^. dsHLA + NV9,^2^ MIIA and GFP-Y66oNBY^3^ protein complexes were expressed recombinantly and purified as described previously and used without further purification. The helix-turn-helix (HTH) peptide has the sequence Ac-AAAAAAAAAAAAAAKYGGGAAAAAAAAAAAAAAK-OH and was purchased from Peptides & Elephants (Hennigsdorf, Germany). Ammonium acetate used was of trace metals basis quality (99.99%, Honeywell), cesium iodide was trace metals basis (99.999%, Sigma-Aldrich), water, trifluoroacetic acid and methanol were LC-MS grade (Merck). Ammonia solution (SupraPur, Supelco) and acetic acid (glacial, Sigma-Aldrich) were used to adjust pH of ammonium acetate solution.

**Instrument modification for access of X-ray light at light source facilities:**

The instrumental setup used for the analysis was based on a high-mass modified Q-ToF2 (quadrupole time-of-flight) mass spectrometer (MS Vision / Micromass) equipped with a quadrupole capable of mass selection up to 30,000 *m/z*.^4^ For coupling with X-ray light sources, the instrument was additionally further modified. Namely, a DN100ISO-K flange was manufactured into the side of the analyzer vacuum chamber at the level of the rf-guiding transfer hexapole between the collision cell and the entrance into the ToF analyzer. This flange was via a custom DN100ISO-K / DN40CF reducer, vacuum isolation valve, a set of short flexible steel bellows and a differential pumping unit mounted onto the open port of the P04 soft X-ray beamline at the PETRA III synchrotron. Through this assembly, X-ray optical access was enabled into the transfer hexapole. There, passing between the hexapole rods, the X-ray light was crossed perpendicularly with the continuous ion beam inside the instrument to enable ion-light interactions. Past this interaction zone, the X-ray light was visualized on a 20×0.1 mm planar crystal screen of cerium-doped yttrium aluminum garnet (Ce:YAG, SKB10B, CRYTUR Ltd.). This screen was directly observed through two optical viewports located above the transfer hexapole region in the top lid of the analyzer chamber and directly behind the Ce:YAG screen on-axis with the X-ray beam.

For measurements at the FL24 beamline of the FLASH2 X-ray free electron laser facility (also at DESY in Hamburg, Germany), the above setup was slightly modified as follows. First, to protect the Ce:YAG screen as well as the on-axis optical viewport from very high intensity FEL X-ray pulses, a beamstop was fashioned from a block of pure copper, which was mounted on a rotating vacuum manipulator between the transfer hexapole and the Ce:YAG screen. For direct observation of the position of the X-ray beam on the screen, the beamstop was rotated out of the beam path, while the FLASH2 FEL operation was requested in single pulse per 10 Hz pulse train mode and the X-ray light was further attenuated by 2×405 nm thick niobium transmission filters. Further, as FEL operation at FLASH2 is inherently pulsed in contrast to the quasi-continuous synchrotron light at PETRA III, the inert gas-filled collision cell of the mass spectrometer, which was also optionally used to impart additional collision activation to the ions prior to their irradiation in both PETRA III and FLASH2 experiments, was repurposed as a linear ion trap. This allowed accumulating ions and their release in time to coincide inside the transfer hexapole with the arrival of FEL photon pulses. This was achieved by using a TGP110 delayed pulse generator (Thurlby Thandar Instruments) to modulate the DC voltage applied to the exit lens of the collision cell (Fig. 1). While in normal operation this element would be kept at constant -10 V to aid in the extraction of thermalized protein ions from the collision cell, we instead applied +10 V blocking potential on this lens to temporarily stop the ions inside the collision cell. Together with relatively high pressure of inert argon (~ 1.2×10^-2^ mbar) inside the cell, tapered electrodes attracting ions towards the exit lens and radiofrequency confining fields applied to the internal hexapolar rod electrodes of the cell, this effectively turned the device into a linear ion trap and led to the accumulation of thermalized ions close to its exit lens. Subsequently, external trigger signal provided by the FLASH2 facility at a precisely defined time prior to the arrival of a photon pulse train functioned as an external trigger for the TGP110 pulse generator. It then over 5 ms inverted the polarity on the collision cell exit lens to -10 V, effectively ejecting the ions in an ion packet. The exact timing of the ejection was experimentally timed to coincide with the arrival of photons into the interaction zone.

**Methods and parameters:**

Prior to native MS, all proteins were buffer-exchanged into ammonium acetate, which is a common MS-compatible water-based buffer surrogate. This was achieved using either spin gel filtration columns (Zeba-Spin, Thermo Scientific or MicroBio-Spin P6, Bio-Rad) or microconcentrators (Amicon Ultra-0.5, 10 kDa MWCO, Merck Millipore). Proteins in ammonium acetate were diluted to appropriate concentration as listed in Table S1 and admitted into the experimental setup using static nano-ESI from gold-coated in-house pulled borosilicate glass capillaries (Kwik-Fil 1B120F-4, World Precision Instruments) essentially as described before.^5^

For the IM-MS experiments, HTH peptides were dissolved in the ratio of 1 mg of peptide per 1 mL of TFA and 0.1 mL of Water. The solution was introduced into the mass spectrometer with standard ESI with a flow rate of 2-5 μL/minute.

**Table S1.** concentration of protein and the concentration and pH of ammonium acetate solutions used for native MS

|  | PETRA III P04 | | | | FLASH II FL24 |
| --- | --- | --- | --- | --- | --- |
|  | GroEL | ds-HLA | Hb | Mutant-GFP | Hb |
| Sample concentration (μM) | 1 | 17 | 8.5 | 5 | 6 |
| AmAc concentration (mM) | 100 | 300 | 100 | 50 | 100 |
| AmAc pH | 7 | 8 | 8 | 8 | 8 |

**Table S2.** Q-ToF and Photon parameters for the experiments

|  |  | PETRA III P04 | | | | | | FLASH II FL24 |
| --- | --- | --- | --- | --- | --- | --- | --- | --- |
|  |  | GroEL | ds-HLA | | Hb | Mutant-GFP | HTH | Hb |
| QToF ultima prameters | Source pressure (mbar) | 10 | 10 | | 9.5 | 9.4 | 9 | 10 |
|  | CC pressure (mbar, Ar) | 1.70E-02 | 1.80E-02 | | 1.20E-02 | 1.20E-02 | 9.00E-03^#^ | 1.40E-02 |
|  | Capillary voltage (kV) | 1.3 | 1.45 | | 1.2 | 1.3 | 3 | 1.3 |
|  | Sampling cone (V) | 150 | 150 | | 80 | 80 | 120 | 80 |
|  | Pusher interval (µs) | 200 | 200 | | 220 | 220 | 20 | 130 |
|  | LM and HM res | 5 | 5 | | 7 | 7 | 10 | 3 |
|  | precursor m/z | 11942 | 3362 | | 4033 | 3955 | - | 4041 |
|  | CE (V) | 50 | 20^ | 120^ | 10 | 30 | 10 | 50 |
| Photon parameters | Photon energy (eV) | 595* | | | 700 | | 310 | 164 |
|  | Average pulse energy (µJ) | - | | | - | - | - | 140 |

^ collision voltage when the ds-HLA ions are not collisionally activated is 20 V while the activated ones are at 120 V

* Pink (polychromatic) beam

\# Nitrogen used for the HTH ion mobility experiment

**Data acquisition**

The actual data acquisition was also performed differently during PETRA III and FLASH2 experiments. As the PETRA III synchrotron light source could be considered continuous for our purposes, irradiated ions were monitored in all individual pusher pulses into the ToF analyzer and could therefore be detected by the mass spectrometer’s multichannel plate (MCP) detector and processed normally through its 1 GS/s (gigasample per second) time-to-digital converter circuitry. Five thousand such individual ToF measurements were then summed together every second, converted from time domain into mass calibrated *m/z* domain and displayed in Waters MassLynx 4.1 software. At FLASH2, however, due to the short duration of arriving X-ray pulse trains interspersed with >99.8% of ‘dark time’ with no photons, external detection and digitization was used instead. For this, we instead acquired raw pre-amplified ToF signal using a fast oscilloscope (Tektronix MSO70804C) directly coming from the MCP detector. The 8 GHz bandwidth oscilloscope acquired data with constant sampling rate of 3.13 GS/s for 1 ms period at each triggered acquisition, covering about 5 consecutive pusher pulses – individual ToF measurement cycles. Synchronization of the start of the acquisition with the arrival of FEL X-ray pulses as well as with the release of trapped ion release was achieved by externally triggering the oscilloscope acquisition with a separate dedicated trigger signal provided by FLASH2. Relative synchronization of the events was achieved through offset adjustments of the precisely timed trigger signals.

**Data processing**

The raw oscilloscope data were later offline processed in a purpose-written MATLAB script. Due to the impracticality of also synchronizing ToF pusher pulsing, the script first detected the rising edge of the first pusher event showing in every MCP readout as a double peak of sharp electronic spikes. Then, based on the first pusher peak in the acquisition, it aligned all the oscilloscope data traces time-wise and summed them. The relative scarcity of oscilloscope data in contrast to standard continuous acquisition caused by FEL pulse dark time was partially mitigated by employing true analog-to-digital conversion mode, where the intensity and shape of electronic signal peaks in the MCP record was retained in contrast to standard TDC acquisition mode, which only builds signal histograms through counting events surpassing a predefined threshold. This was enabled by higher sampling rate of the used oscilloscope digitizer in contrast to the original TDC card inside the Q-ToF2 electronics and thus better peak shape retention due to denser signal sampling sufficient to capture the peak shape. Time-aligned and summed raw data were then split into individual pushes / ToF measurements and based on later evaluation, second up to four pushes from each trace, which contained signals of arriving ions, were further summed. Hence overall, 3×5000 individual ToF measurements were always summed per each final mass spectrum. Prior to data analysis, such raw spectra in the ToF domain were converted into the *m/z* domain using the standard ToF equation (Eq. 1)

**Equation 1:**

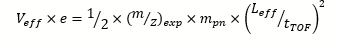

where *V_eff_* is the effective acceleration voltage experienced by ions entering the ToF, *e* is the elementary charge (1.60×10^-19^ C), *m_pn_* is the approximate mass of proton or neutron (1.67×10^-27^ kg or 1.67×10^-24^ g), *L_eff_* is the effective length of the ToF trajectory, *t_TOF_* is the experimentally measured time-of-flight of an ion and finally where *(m/z)_exp_* is the resulting mass-over-charge ratio of the ion reflecting merely experimental instrument’s geometry without mass recalibration.

The resulting mass spectra were then mass recalibrated using the time-of-flight *t_0_* defined by the rising edge of the electronic pusher peak and utilizing a set of fifth-order polynomial coefficients derived from a measured spectrum of known mass clusters of caesium iodide through equation 2.

**Equation 2:**

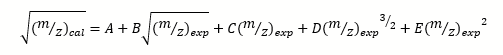

where *(m/z)_cal_* and *(m/z)_exp_* are recalibrated and raw *m/z* of an ion, respectively, and *A* to *E* are experimentally derived calibration coefficients obtained from a spectrum of known CsI clusters. Finally, both synchrotron data processed with MassLynx as well as FEL data processed manually as described above were read into a custom Python 3.11 script and plotted using the standard matplotlib 3.8.2 library.
